# Cell culture system to efficiently test candidate genes and molecular pathways implicated in congenital diaphragmatic hernias

**DOI:** 10.1101/2020.04.03.024851

**Authors:** Eric L. Bogenschutz, Elizabeth M. Sefton, Gabrielle Kardon

**Affiliations:** Department of Human Genetics, University of Utah, Salt Lake City, UT 84112

**Keywords:** CDH, fibroblasts, diaphragm, congenital diaphragmatic hernia, cell culture

## Abstract

The mammalian muscularized diaphragm is essential for respiration and defects in the developing diaphragm cause a common and frequently lethal birth defect, congenital diaphragmatic hernia (CDH). Human genetic studies have implicated more than 150 genes and multiple molecular pathways in CDH, but few of these have been validated because of the expense and time to generate mouse mutants. The pleuroperitoneal folds (PPFs) are transient embryonic structures in diaphragm development and a critical cellular source of CDH. We have developed a system to culture PPF fibroblasts from E12.5 mouse embryos and show that these fibroblasts, in contrast to the commonly used NIH 3T3 fibroblasts, maintain expression of key genes in normal diaphragm development. Using pharmacological and genetic manipulations that result in CDH *in vivo*, we also demonstrate that differences in proliferation provide a rapid means of distinguishing healthy and impaired PPF fibroblasts. Thus, the PPF fibroblast cell culture system is an efficient tool for testing the functional significance of CDH candidate genes and molecular pathways and will be an important resource for elucidating the complex etiology of CDH.

**SUMMARY STATEMENT:** Multiple candidate genes and molecular pathways have been identified as potentially contributing to the etiology of congenital diaphragmatic hernias. We describe a cell culture system to efficiently functionally test these candidates.

## INTRODUCTION

The diaphragm is a mammalian-specific skeletal muscle that is essential for the inspiration phase of respiration and separates the thoracic from the abdominal cavity. Defects in diaphragm development are relatively common, occurring in 1 in 3000 births (Stege et al., 2003). In congenital diaphragmatic hernias (CDHs), abdominal contents herniate through weakened portions of the diaphragm into the thoracic cavity. As a result, lung development is impeded, leading to lung hypoplasia and high neonatal mortality (Pober, 2007). Genetic studies in humans have identified multiple chromosomal abnormalities and *de novo* mutations in numerous genes that may contribute to the etiology of CDH and have implicated the involvement of multiple molecular pathways (reviewed by Kardon et al., 2017). However functional validation that mutations in any of these candidate genes are true causal variants or demonstration that a particular molecular pathway is critical for development of CDH has been a significant challenge.

The diaphragm develops primarily from two embryonic tissues (Merrell et al., 2015). The pleuroperitoneal folds (PPFs) are transient bilateral pyramidal structures derived from lateral plate mesoderm and located between the thoracic (pleural) and abdominal (peritoneal) cavities (Merrell et al., 2015) and ultimately give rise to the diaphragm’s muscle connective tissue and central tendon. The somites are the source of migratory muscle progenitors which target the PPFs and subsequently give rise to the diaphragm’s muscle (Allan and Greer, 1997; Dietrich et al., 1999; Babiuk et al., 2003; Sefton et al., 2018). As the muscle progenitors enter the PPFs, the PPFs spread dorsally and ventrally to form a continuous sheet in which the central tendon differentiates in the central region and the muscle differentiates peripherally to form the radial array of costal myofibers (Merrell et al., 2015; Sefton et al., 2018). Importantly, the development of the PPFs has been found to regulate the overall morphogenesis of the diaphragm (Babiuk et al., 2003; Merrell et al., 2015; Sefton et al., 2018).

Recent studies have established that the PPFs not only control normal diaphragm development, but are a critical cellular source of CDH. Several mouse conditional mutagenesis studies have demonstrated that CDH-implicated genes cause hernias when mutated in PPFs or their associated mesothelium (Merrell et al., 2015; Paris et al., 2015; Carmona et al., 2016). In addition, research from our lab on the transcription factor *Gata4*, a gene strongly implicated in CDH by human genetic studies (Arrington et al., 2012; Longoni et al., 2012; Wat et al., 2012; Yu et al., 2012; Longoni et al., 2014; Kammoun et al., 2018), has specifically demonstrated that deletion of *Gata4* in the PPF fibroblasts, and not muscle progenitors, leads to CDH with complete penetrance (Merrell et al., 2015). While deletion of CDH-implicated genes leads to herniation beginning at E16.5 in mouse (E60 in humans), these genes are required much earlier in the PPFs – between E10.5 and 12.5 (E28 - E40 in humans; Merrell et al., 2015; Paris et al., 2015; Carmona et al., 2016) and lead to both cell-autonomous changes within the PPF fibroblasts and cell non-autonomous effects on muscle progenitors. With deletion of CDH genes, PPF fibroblasts exhibit decreased proliferation and increased apoptosis as well as altered gene expression (Merrell et al., 2015; Paris et al., 2015; Carmona et al., 2016) and also affect the proliferation of neighboring muscle progenitors to ultimately lead to weakened regions of the diaphragm that allow herniation (Merrell et al., 2015). Altogether these *in vivo* studies have established that changes in the proliferation, survival, and gene expression of PPFs between E10.5 and E12.5 are critical determinants of whether CDH develops.

Besides mutations in individual genes, alterations in molecular signaling pathways regulating diaphragm development have been implicated in CDH (Kardon et al., 2017). Multiple pathways, including FGF, Hedgehog, and Wnt/β-catenin signaling have been implicated. Most notably, alterations in maternal vitamin A or its derivative retinoic acid (RA) in the embryo have been proposed to be a potent source of CDH (Clugston et al., 2010a). RA is synthesized from vitamin A (retinol) via a two-step oxidation process: alcohol or retinol dehydrogenases convert retinol to all-trans-retinal and retinaldehyde dehydrogenases (primarily ALDH1A2 – formerly known as RALDH2 in the embryo) converts ATRAL to all-trans-retinoic acid (RA or ATRA) (Shannon et al., 2017). RA functions as either a paracrine or autocrine signal, binding retinoic acid receptors to form heterodimer complexes with retinoid receptors and activate transcription (Rhinn and Dollé, 2012; Cunningham and Duester, 2015; Shannon et al., 2017). The importance of RA signaling in CDH was first demonstrated in rodents where low maternal vitamin A (Anderson, 1941; Wilson et al., 1953) or environmental teratogens that interfere with RA signaling induces diaphragmatic hernias in offspring (Mey et al., 2003; Babiuk et al., 2004; Noble et al., 2007; Clugston et al., 2010a; Clugston et al., 2010b). In addition, babies born with CDH have been found to have low retinol and retinol binding protein in their blood (Major et al., 1998; Beurskens et al., 2010). Finally, mutations in retinoic acid receptors in mice and humans have been associated with CDH (Mendelsohn et al., 1994; You et al., 2005; Golzio et al., 2007; Pasutto et al., 2007). Thus, both environmental perturbations of and genetic mutations in RA signaling have been implicated in CDH. Mechanistic insights into how disruptions in RA signaling cause CDH have largely come from delivery of teratogens to pregnant rodents. Nitrofen is a diphenyl ether herbicide that inhibits ALDH1A2 activity (Mey et al., 2003), and its delivery induces diaphragmatic hernias in embryos (Costlow and Manson, 1981; Mey et al., 2003; Babiuk et al., 2004; Noble et al., 2007; See et al., 2008; Clugston et al., 2010a; Clugston et al., 2010b). ALDH1A2 is strongly expressed in the PPFs (Mey et al., 2003; Clugston et al., 2010a), and nitrofen inhibits proliferation of PPF fibroblasts and overall PPF morphogenesis (Clugston et al., 2010b). These teratogen studies demonstrate that, like mutations in individual CDH-implicated genes, inhibition of RA signaling induces CDH via its inhibition of PPF growth and morphogenesis.

Overall, the etiology of CDH is complex and heterogeneous, potentially involving more than 150 genes and multiple molecular pathways, but few of these genes and pathways have been functionally validated in mice as causative of diaphragmatic hernias (Kardon et al., 2017). This is in part due to the cost and time required to generate and analyze suitable mouse models. As the diaphragm is a mammalian-specific structure (Perry et al., 2010), rodents are the only widely used animal model to study CDH *in vivo*. Rodent *in vivo* studies indicate that the PPFs are critical for diaphragm development, and inhibition of early PPF growth and morphogenesis, via genetic mutations or inhibition of molecular signaling pathways, leads to CDH. Here, we develop a system to culture these early PPFs and show they maintain expression of key genes in normal diaphragm development. Furthermore, we validate that genetic and pharmacological manipulations known to cause hernias *in vivo* in mice lead to impaired growth of PPF fibroblasts *in vitro*. This cell culture system, in combination with pharmacological interventions and future gene knock-down and editing experiments, will allow for rapid screening of CDH candidate genes and molecular pathways and provide an important tool for prioritizing genes to functionally test *in vivo* in mice and ultimately target therapeutically.

## RESULTS AND DISCUSSION

### Early pleuroperitoneal folds can be isolated and expanded in culture

PPF fibroblasts are critical in both diaphragm development and an important cellular source of CDH. Because defects in proliferation and the morphogenetic spread of early PPF fibroblasts are essential for the development of CDH *in vivo* (Clugston et al., 2010b; Merrell and Kardon, 2013; Paris et al., 2015; Carmona et al., 2016), we endeavored to set up an *in vitro* system to determine whether E11.5-E12.5 PPF fibroblasts can be cultured and maintain their ability to proliferate and spread. PPFs at this stage are bilateral pyramidal structures composed predominantly of fibroblasts and a smaller number of muscle progenitors which have migrated into the PPFs (Sefton et al., 2018). The PPFs are visibly distinct by E11.5, and E11.5 - E12.5 PPFs are able to be manually dissected from the underlying liver with forceps under a dissecting microscope (**Fig. 1A, Video 1**). Isolated pairs of PPFs are removed with sterile forceps and then plated whole, ventral-side down, on tissue-culture treated 96 well plates with PPF growth media (F12 media, 10% FBS and 50 μg/ml Gentamicin). Once plated, the PPFs proliferate and spread across the well, similar to PPFs *in vivo* (**Fig. 1B**; **Video 2**). While E11.5 – E12.5 PPFs initially contain muscle progenitors, we found that these growth conditions do not favor myogenic cells and muscle progenitors, which express Pax7, are neither present (**Fig. S1**) nor differentiate into myotubes after 5 days of culture. Thus these conditions allow us to specifically assess the proliferation and spread of the PPF fibroblasts.

**Figure 1.**
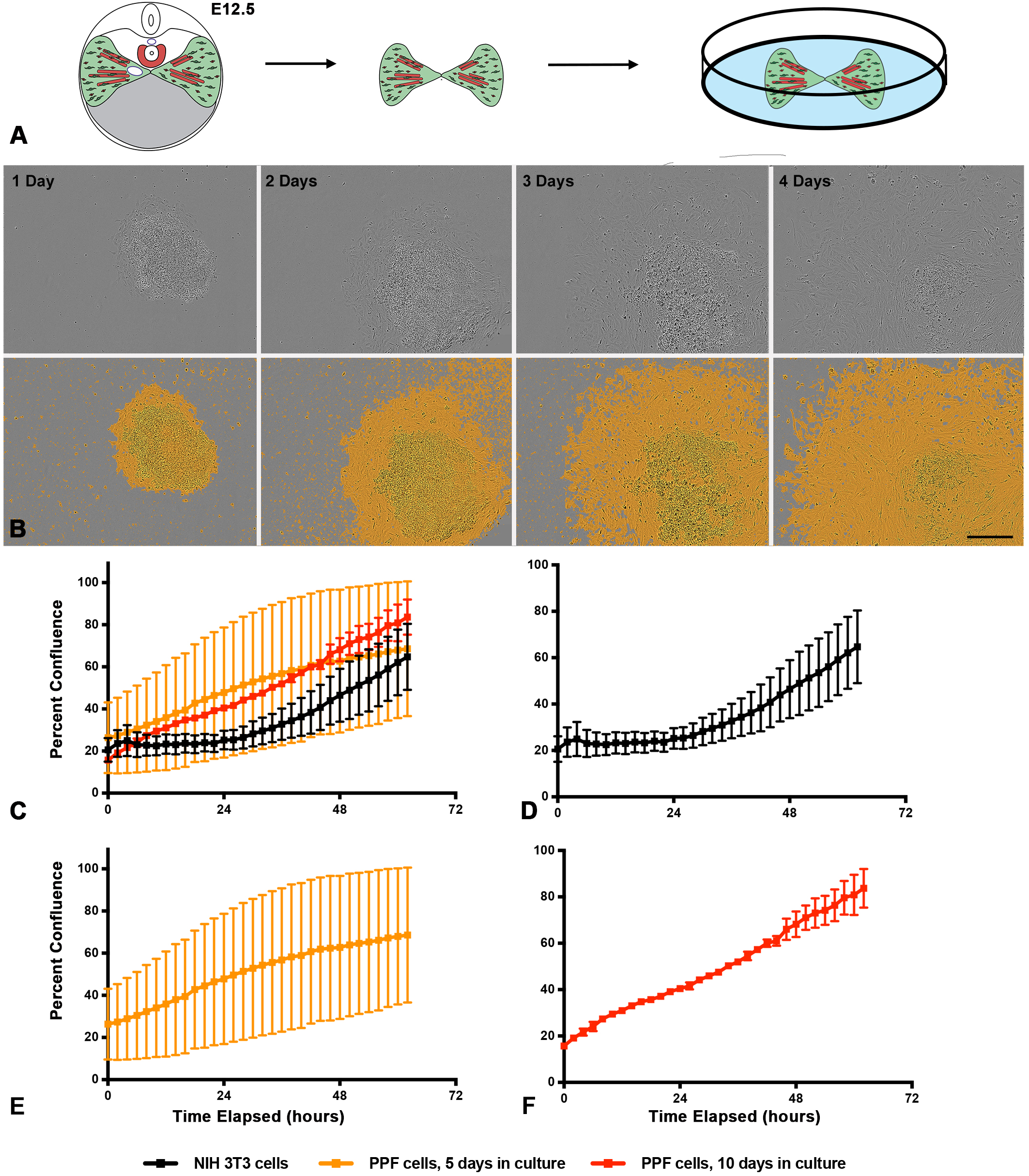
PPFs can be isolated and cultured *in vitro*. **A**. E12.5 pairs of PPFs, including muscle progenitors, are dissected from embryos and cultured individually in single wells of 96-well culture plates. **B**. Growth of E11.5 PPFs 1 - 4 d after isolated. Top row, phase-contrast images taken by IncuCyte cell imager. Bottom row, images with IncuCyte created confluency mask layer to highlight cells (in orange). Scale bar is 300 μm. **C-F**. Comparison of growth of PPFs and NIH3T3 fibroblast (**C**). Growth of NIH 3T3 cells (**D**, n = 9 biological replicates), PPF cells after 5 d in culture and 1 passage (**E**, n = 7 biological replicates included, 4 poor growth wells excluded) and PPF cells after 10 d in culture and 2 passages (**F**, n = 2 biological replicates included, 2 poor growth wells excluded). Growth is measured by percent confluency calculated by IncuCyte software and for each timepoint the average of the biological replicates are shown with error bars representing standard error of the mean (SEM).

Our culture conditions allow E12.5 PPF fibroblasts to successfully grow up to 14 d culture. During the initial culture of the intact PPFs, cells expand radially, but after 5 d in culture PPF cells become highly confluent and stop growing; at this point PPF fibroblasts are dissociated from each well, disaggregated, resuspended, and re-plated at an even density in a new well. The PPF fibroblasts can then be grown an additional 5 d, passaged, splitting cells from 1 well into 2-3 wells (with ∼10,000 cells seeded at an even density in each well) and grown for a final 3-4 d. After the first passage at 5d or after the second passage at a total of 10 d in culture, we imaged and monitored cell proliferation for 62 hours using a Sartorius IncuCyte ZOOM live cell imager (**Fig. 1C-F, Fig. S2**). In addition, we compared the growth of PPF fibroblasts with NIH 3T3 fibroblast cells, immortalized fibroblasts derived from Swiss mouse embryonic fibroblasts (Todaro and Green, 1963), that were grown and imaged on the same 96 well plate (**Fig. 1D, Fig. S2B**). As expected, the NIH 3T3 fibroblasts grow under these conditions with little variability and double every 38 h (+/- 5 h, **Fig. 1D, Fig. S2B**). The PPFs grow more variably. After the first and second passages, the number of PPF fibroblasts that adhere to the wells can be variable; wells with higher initial density of cells reliably grow better in comparison to those with low seeding density (**Fig. S2C-D**). Excluding the obvious wells that did not grow well, we find that PPF fibroblasts after 5 d in culture double every 39 h (+/- 9.6 h) and after 10 d in culture every 23 h (+/- 1.7 h, **Fig. 1 C-D**). Thus, we demonstrate that PPFs can be successfully isolated, PPF fibroblasts grown in culture and, with careful attention to the initial density of PPF fibroblasts, the growth rate of these cells determined.

Maintaining muscle progenitors in combination with PPF fibroblasts may be important for assaying phenotypes associated with the interaction between myogenic cells and fibroblasts. PPFs grown in PPF growth media gradually lose any Pax7+ myogenic progenitors present in the initial PPF explant (**Fig. S1**). To optimize conditions for myogenic cells, we cultured PPFs from *Pax3*^*Cre/+*^; *Rosa*^*nTnG/+*^ mice. *Pax3*^*Cre*^ recombines in the somites and earliest muscle progenitors (Engleka et al., 2005) and in combination with the Cre-responsive reporter *Rosa*^*nTnG*^ (in which all cells express nuclear Tomato and in response to Cre cells express nuclear GFP) labels all myogenic cells with GFP and PPF fibroblasts with Tomato. E12.5 PPFs from *Pax3*^*Cre/+*^; *Rosa*^*nTnG/+*^ mice were cultured with 0.5 nM FGF (Farina et al., 2012) and 10% fetal bovine serum in either F-10 or F-12/DMEM media (1:1) to compare muscle progenitor survival over time. Over 5 d in culture, while F-10 did not robustly support GFP+ myogenic cells (**Fig. 2A-B**), F-12/DMEM cultures showed increased numbers of GFP+ myogenic cells (**Fig. 2C-D**). Thus, for analyses requiring co-culture of myogenic cells and PPF fibroblasts, F-12/DMEM media with 10% FBS and FGF is a suitable media.

**Figure 2.**
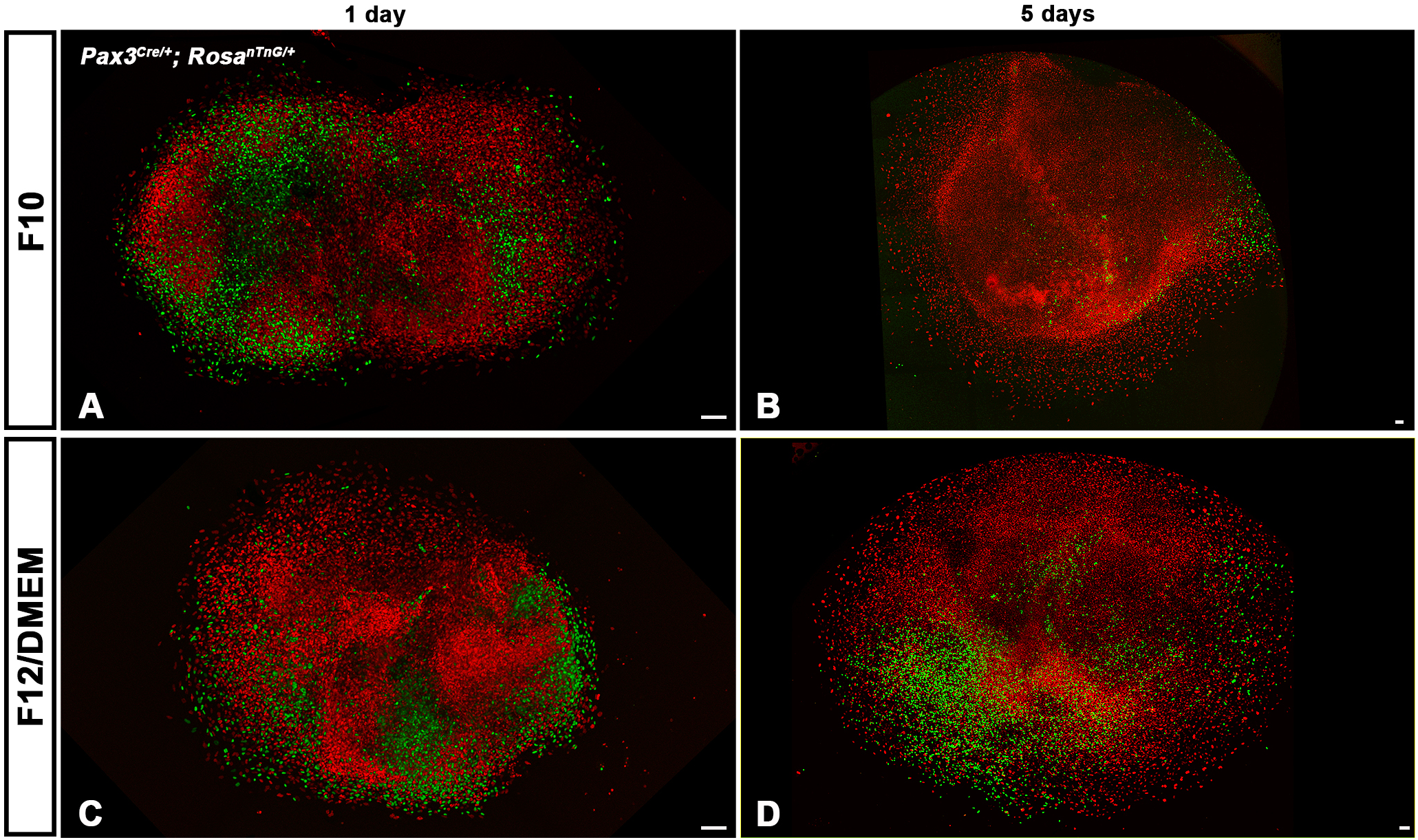
Muscle progenitors and fibroblasts cultured in myogenic media. *Pax3*^*CreKI/+*^; *Rosa*^*nTnG/+*^ E12.5 PPFs, cultured for 1 d (**A, C**) or 5 d (**B, D**) in F10 with FBS and FGF (**A, B**) or F12/DMEM with FBS and FGF (**C, D**). F10 media is only suitable to expand PPF fibroblasts, as indicated by the predominance of Tomato+ PPF fibroblasts and lack of GFP+ myogenic cells (n = 3) (**B**). F12/DMEM allows expansion of both GFP+ muscle progenitors and Tomato+ PPF fibroblasts as large numbers of GFP+ cells were found in 3 of 4 PPF cultures in this media (**D**). Scale bars are 100 μm.

### E12.5 PPF fibroblasts maintain expression of key diaphragm genes for at least 10 days in culture

A robust PPF cell culture model of early diaphragm development and CDH requires that expression of key genes important for normal diaphragm development (and aberrant in CDH) is maintained in wild-type PPF fibroblasts during culture. As such, we examined the expression of multiple diaphragm genes during culture of E12.5 PPF fibroblasts. We included the transcription factors *Gata4* (Jay et al., 2007; Yu et al., 2012; Merrell et al., 2015), *Gata6* (Yu et al., 2014), and *Tbx5* (which can bind cooperatively with Gata4 to transactivate genes; Valasek et al., 2011; Ang et al., 2016); Gata4 co-factor *Zfpm2* (Ackerman et al., 2005; Chlon and Crispino, 2012); and the orphan nuclear receptor *Nr2f2* (which binds and interacts with *Gata4* and *Zfpm2*; Huggins et al., 2001; You et al., 2005). Together the proteins encoded by these genes likely form an essential transcriptional network regulating normal diaphragm development and each one of these genes when mutated causes CDH. We also included the receptor *Pdgfr*α which labels muscle connective tissue fibroblasts and is implicated in CDH (Bleyl et al., 2007; Uezumi et al., 2010). We found that all of these genes are expressed in E12.5 PPFs at harvest and *Gata4, Zfpm2, Nr2f2, Tbx5*, and *Gata6* do not change significantly after 5 or 10 d in culture (**Fig. 3A-E**). Only *Pdgfra* is somewhat altered during culture, as it is transiently downregulated after 5 d in culture but recovers expression following 10 d in culture (**Fig. 3F**). In contrast, while NIH 3T3 fibroblasts express similar levels of *Gata4, Gata6, Nr2f2*, and *Pdgfra* as E12.5 PPFs at harvest, they expressed significantly lower levels of *Zfpm2* and undetectable levels *Tbx5* (**Fig. 3A-F**). As regulation of Gata4 transcriptional activity by Zfpm2 and Tbx5 is essential for regulating normal diaphragm development, this suggests that the transcriptional activity of NIH 3T3 cells will differ significantly from that of PPF fibroblasts. Thus, these experiments show that E12.5 PPF fibroblasts, and not NIH 3T3 fibroblasts, largely maintain expression of a network of key diaphragm genes after 10 d in culture.

**Figure 3.**
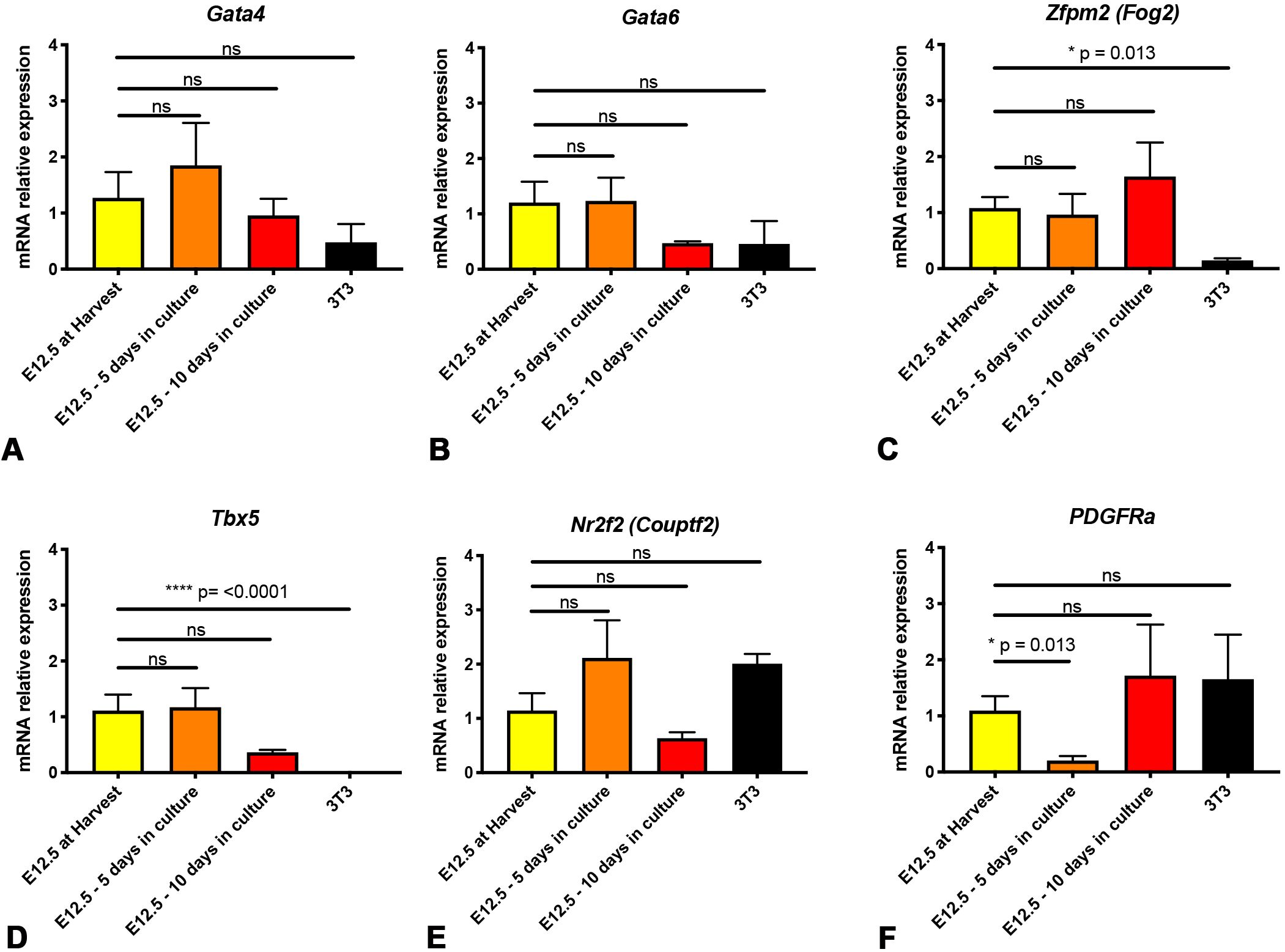
Expression of important diaphragm genes is maintained in E12.5 PPFs during culture. Expression of *Gata4* (**A**), *Zfpm2* (**B**), *Nr2f2* (**C**), *Tbx5* (**D**), and *Gata6* (**E**) does not statistically differ in E12.5 PPFs when cultured 5 or 10 d. *PDGFRa* is reduced after 5 d in culture, but is not significantly reduced after 10 d in culture (**F**). NIH 3T3 fibroblasts have lower expression of *Zfpm2* (**C**) and undetectable levels of *Tbx5* (**D**). Gene expression determined by qRT-PCR, normalized against *18S* gene expression, and then normalized to “at harvest” value, which was set to 1. Significance tested with one-way ANOVA. Holm–Sidak post hoc tests are indicated by * p<0.05, **** p<0.0001. Error bars represent standard error of the mean (SEM).

As early developmental events are critical to the development of CDH (Merrell et al., 2015; Paris et al., 2015; Carmona et al., 2016), we also analyzed gene expression in cultured E11.5 PPFs (**Fig. S3**). First, we compared E11.5 to E12.5 PPFs at harvest and found no significant differences in expression of the important diaphragm genes (**Fig. S2G**). However after 5 or 10 d in culture, while expression of *Gata4* and *Gata6* are maintained (**Fig. S3A-B**), *Zfpm2, Tbx5, Nr2f2*, and *PDGFRa* are significantly reduced in E11.5 PPF fibroblasts (**Fig. S3C-F**). The reduced expression of multiple critical genes during culture of E11.5 PPFs indicates that these younger PPFs are not adequate as a cell culture model. Instead E12.5 PPF fibroblasts, but not E11.5 PPF or NIH 3T3 fibroblasts, continue to express important diaphragm genes in culture and likely serve as a good *in vitro* system for studying development of the diaphragm and CDH.

### Genetic and pharmacological manipulations that lead to CDH in vivo cause decreased PPF proliferation in vitro

Rodent *in vivo* studies demonstrate that inhibition of early PPF growth and spread, via genetic mutations or pharmacological inhibition of molecular signaling pathways, leads to CDH (Clugston et al., 2010b; Merrell et al., 2015; Paris et al., 2015) and suggest that reduction in the proliferation of early PPF cells may be a common feature of CDH etiology. Therefore, we sought to determine whether a genetic mutation and pharmacological inhibition of a signaling pathway known to cause CDH *in vivo* would be recapitulated as changes in E12.5 PPF proliferation *in vitro*.

Mutations in *Gata4* have been strongly implicated by human genetic studies to cause CDH (e.g. Yu et al., 2013), and our lab has demonstrated in mice *in vivo* that loss of *Gata4* in PPF cells leads to decreased proliferation of PPF cells at E12.5 and CDH after E16.5 (Merrell et al., 2015). To test whether genetic loss of *Gata4* leads to decreased proliferation of PPFs *in vitro*, we harvested E12.5 PPFs from embryos in which either one (*Prx1Cre*^*Tg/+*^;*Gata4*^*fl/+*^;*Rosa*^*mTmG/+*^) or both alleles (*Prx1Cre*^*Tg/+*^;*Gata4*^*del/fl*^;*Rosa*^*mTmG/+*^) of *Gata4* were deleted with the *Prx1Cre* transgene that is expressed in PPFs (Logan et al., 2002). As previously noted (Merrell et al., 2015), the *Prx1Cre* transgene does not recombine in all PPF cells by E12.5. Thus the Cre-responsive *Rosa*^*mTmG*^ reporter (Muzumdar et al., 2007) was used to identify GFP+ PPF fibroblasts in which the Cre protein was expressed and either one or two *Gata4* alleles was deleted. In order to determine the effects of *Gata4* deletion on proliferation of PPF fibroblasts, we analyzed growth of all PPFs (with phase images) and growth of GFP+ PPFs (with GFP fluorescence images) using the IncuCyte ZOOM live cell imager. We compared the fold change in the GFP+ PPF fibroblasts (% confluence at time T/% confluence at time 0) in which one or two alleles of *Gata4* was deleted with the fold change in all PPF fibroblasts and plotted this ratio over time. We found that the proportion of GFP+ PPF fold change relative to the total cell fold change decreased with time when both alleles of GFP were deleted (red line versus blue line in **Fig. 4A**). By repeated measures ANOVA, this difference is statistically significant when comparing proliferation during 48-90 hours (**Fig. 4A**). These data indicate that loss of the important CDH gene *Gata4* inhibits proliferation of E12.5 PPF fibroblasts in culture and suggests that mutations in other genes causing CDH *in vivo* will lead to detectable changes in PPF fibroblasts proliferation.

**Figure 4.**
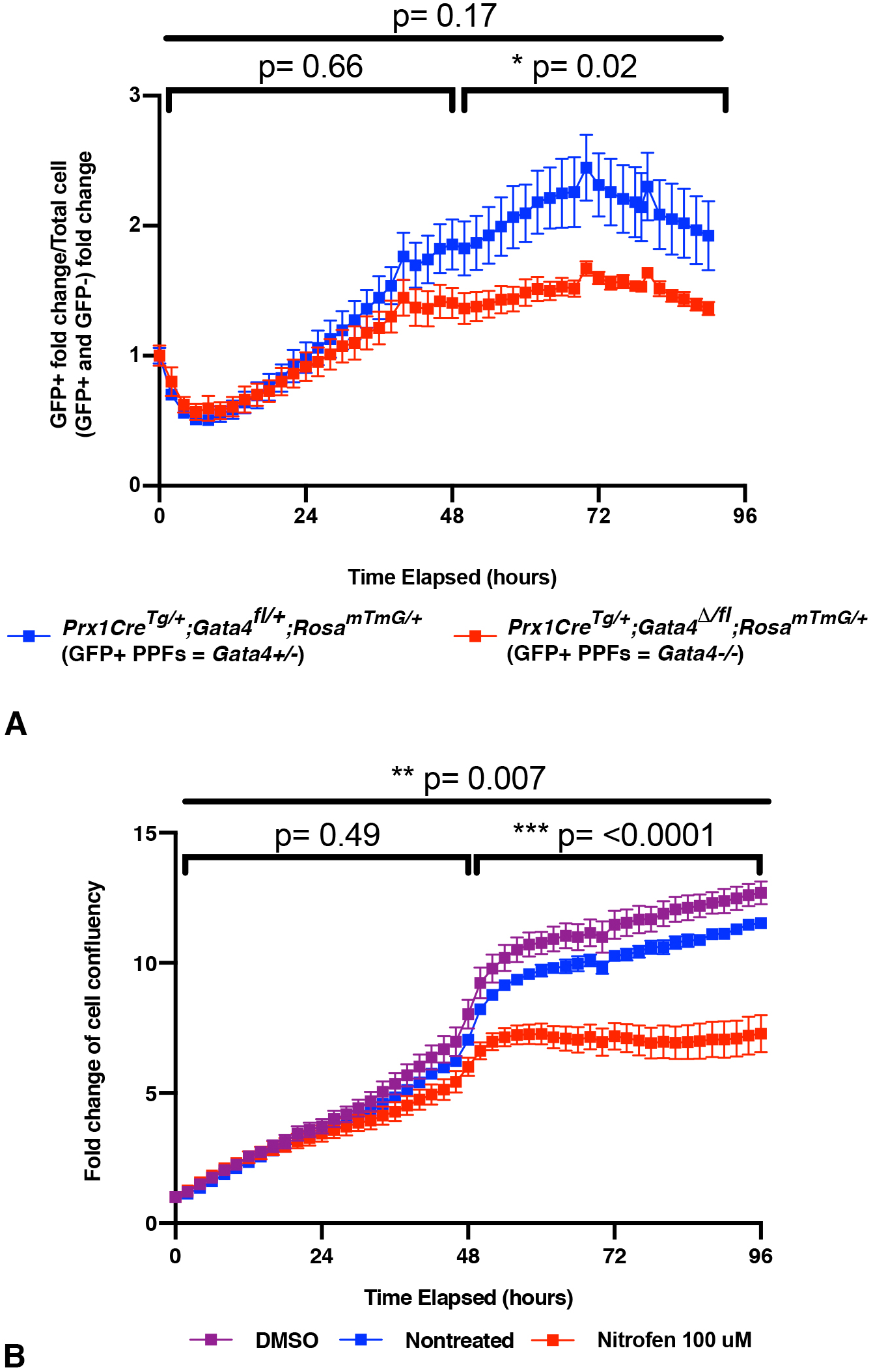
*In vitro* proliferation of PPF fibroblasts is inhibited by genetic mutation and pharmacological inhibitor previously demonstrated to cause CDH *in vivo*. **A**. E12.5 PPFs isolated from either *Prx1Cre*^*Tg/+*^;*Gata4*^*fl/+*^;*Rosa*^*mTmG/+*^ (with GFP+ *Gata4+/-* PPFs) or *Prx1Cre*^*Tg/+*^;*Gata4*^*del/fl*^;*Rosa*^*mTmG/+*^ (with GFP+ *Gata4 -/-* PPFs) embryos and cultured after 5 d (one passage) on the IncuCyte ZOOM, with GFP (to measure confluence of GFP+ cells) and phase (to measure confluence of all cells) images taken. The average ratio GFP+ PPF fold change (% confluence at time T/% confluence at time 0) over total PPF+ fold change +/- SEM is plotted. **B**. E12.5 PPFs were cultured after 5 d (and one passage) either untreated (n = 6), treated with DMSO (n = 6), or treated with nitrofen (n = 6) on the IncuCyte ZOOM to track cell confluency via phase images. Average fold change (% confluence at time T/% confluence at time 0) +/- SEM is plotted. Statistical changes in total confluency over time were determined using repeated measure ANOVA on the log2 transformed fold change over time, with bars denoting whether comparison was between T=0 to T= total time elapsed, or through the first or last halves of the experiments. *** P= <0.001, * P= <0.05.

RA signaling is an important regulator of normal diaphragm development and when inhibited causes CDH (Clugston et al., 2010a). Pharmacological inhibition of RA synthesis via administration to pregnant females of nitrofen, a herbicide that inhibits ALDH1A2 (Alles et al., 1995; Mey et al., 2003; Kling et al., 2010), inhibits growth of the PPFs to ultimately cause CDH in embryos (Costlow and Manson, 1981; Nakao and Ueki, 1987). To test whether nitrofen similarly affects PPF proliferation *in vitro*, we cultured E12.5 PPFs in the presence of 100 µM nitrofen. Compared with untreated or vehicle-treated PPFs, nitrofen treatment significantly decreased PPF proliferation (**Fig. 4B**). Thus the effects of pharmacological inhibition of RA *in vivo* are similarly recapitulated as changes in PPF proliferation *in vitro*.

In summary, these two sets of experiments demonstrate that genetic manipulations and pharmacological inhibitors that cause CDH *in vivo* have a measurable effect on proliferation of E12.5 PPFs when cultured *in* vitro and suggests that this culture system will be an effective tool for functional testing candidate CDH genes and molecular pathways.

### Conclusion

The etiology of CDH is complex and potentially involves mutations in many genes and aberrations in multiple signaling pathways. However, functional testing of candidate genes and molecular pathways for their role in CDH etiology has been limited by the substantial time and expense to generate *in vivo* mouse models. We have developed a more rapid and cost-effective *in vitro* system to carry out functional tests. PPF fibroblasts are a critical cellular source of CDH and we have optimized a protocol to isolate and culture E12.5 mouse PPF fibroblasts, allowing visualization and quantification of the cell population driving normal diaphragm morphogenesis and CDH. In culture, PPF fibroblasts proliferate and maintain expression of key diaphragm development genes. Furthermore, genetic mutations and pharmacological manipulations that lead to CDH *in vivo* impair PPF proliferation *in vitro*. This *in vitro* system, in combination with the large number of pharmacological tools currently available, will enable a wide variety of candidate signaling pathways to be quickly tested and give mechanistic insights into the regulation of PPF growth and development of CDH. In the future, as CRISPR-Cas9 genome editing is optimized for primary cells, mutations in candidate CDH genes can be engineered directly into wild-type PPFs to test their possible function.

## Author Contributions

E.M.S. and E.L.B. performed experiments, data analysis, and drafted the manuscript. E.L. B., E.M.S. and G.K. designed the research. G.K. revised and edited the manuscript.

The authors declare no conflicts of interest.

## MATERIALS AND METHODS

### Cell culture, media, and reagents

NIH/3T3 cells were obtained from ATCC (CRL-1658). Embryos were dissected from pregnant CD-1 female mice mated with CD-1 males or pregnant *Gata4*^*fl/fl*^;*Rosa*^*mTmG/mTmG*^ females mated with *Prx1Cre*^*Tg/+*^;*Gata4*^*del/+*^ males. Staged E11.5 or E12.5 embryos were dissected from yolk sacs in sterile PBS and transferred to Ham’s F-12 Nutrient Mixture (F-12) (Gibco). Embryos were trimmed with two cuts to isolate the trunk region: one cut across the embryo just caudal to the forelimbs and another cut across the embryo just cranial to the hindlimbs (and leaving liver attached to trunk). Developing heart and lungs were pulled out cranially from the thoracic cavity using forceps and the trunk trimmed to the base of the rib cage to expose the PPFs lying lateral to the vertebral column and on top of the liver. Trimmed trunks were pinned onto a Sylgard coated 6mm dish in media. Pairs of PPFs were isolated manually with forceps from the lateral body wall and from the underlying septum transversum and liver (**Video 1**) and then each pair placed in a single well in a 96-well plate (Eppendorf) containing 100 μL of media. To promote growth of PPF fibroblasts, we used PPF growth media: Ham’s F-12 Nutrient Mixture, 10% Fetal Bovine Serum (FBS), 50 μg/mL Gentamycin (all from Gibco). To promote growth of PPF fibroblasts and myogenic cells, we used myogenic media: Ham’s F-10 with L-glutamine (Caisson) or DMEM/F-12 GlutaMAX (Invitrogen), 10% FBS, 50μg/mL gentamycin, and 0.5 nM FGF. PPFs were grown in a 37°C incubator for 5 d, changing media in the wells every 2 d. After 5 d, cells were passaged by removing the media in the well, washing the cells with 100 μl of F-12 media twice, then de-adhering cells by placing 100 μl 0.25% trypsin-EDTA (Gibco) in each well with PPFs and incubating for 9 minutes at 37°C. Trypsin and cells were transferred from each well to a 1.5mL Eppendorf tubes and trypsin neutralized by adding 100 μl of growth or myogenic media. Suspended cells were pelleted by centrifuging for 5 minutes at 340xg, trypsin and media removed, disaggregated cells were resuspended in 100 μl growth or myogenic media and simply transferred to a new well on a 96-well plate. Cells were grown for another 5 d (with media changes every 5 d) and at a total of 10d in culture were passaged again. On this second passage, resuspended cells were counted and ∼10,000 cells/well were seeded and grown for another 3 d. PPF cells were never grown for more than 2 weeks. For nitrofen treatments, nitrofen (Sigma) was dissolved in DMSO for a stock concentration of 100 mM and then diluted 1:1000 (to 100 μM final concentration) in growth media. As a vehicle control, a matching volume of DMSO was also added to growth media. Fresh nitrofen and DMSO were added every 24 hours and media replaced every 2 d.

### Cell growth analysis

Proliferating cells were imaged using the IncuCyte ZOOM which took 4 representative phase-contrast images per well every 2 hours. IncuCyte software was then used to analyze average confluency per well per time point to calculate cell growth over time. To control for differences in seeding densities between wells, fold changes of cell growth were calculated by dividing each treatment by the mean initial confluency at time 0. For GFP experiments, both phase and GFP images were taken by the IncuCyte ZOOM, with the same number of images and time intervals used. Both total cell percentage (from phase-contrast images) and GFP-positive cell percentages (from GFP images) were used to analyze growth of GFP+ cells compared with growth of all cells (from phase-contrast images).

### RNA extraction, cDNA synthesis, and quantitative polymerase chain reaction (qPCR)

Total RNA was extracted with the *Quick*-RNA Microprep kit (Zymo, Irvine, CA) according to manufacturer’s protocol. cDNA was synthesized using Applied Biosystems High-Capacity RNA-to-cDNA kit (ThermoFisher) from purified RNA according to manufacturer’s protocol. qRT-PCR was used to analyze expression of *Gata4, Zfpm2, Nr2f2, Tbx5, Gata6, PDGFRa*, and *Pax7* using pre-validated primer sets (TaqMan, ThermoFisher; **Table S1**). 10 μl or 20 μl reaction volumes were prepared using TaqMan Fast Advanced Master Mix (ThermoFisher). The amplification was performed under the following conditions: 20 seconds at 95°C followed by 40 cycles at 95°C for 1 second and 60°C for 20 seconds. Gene expression levels were normalized against *18S* ribosomal RNA for each sample and fold changes were calculated using 2^-ΔΔCt^ method (Schmittgen and Livak, 2008) by setting the expression levels of each gene at stage-matched harvest as 1. Data collected from 2-5 biological replicates was calculated and plotted as average fold changes with standard error of the mean (SEM).

### Statistical Analysis

Data are presented as mean ± SEM. One-way analysis of variance (ANOVA) was used to determine differences in gene expression. If significant, a Holm-Sidak post hoc test was used. For growth comparisons between either chemical treatments or genotype, repeated ANOVA analysis was run on the log2 fold change of either cell confluence (**Fig. 4A**) or the log2 fold change of GFP cells over total cells (**Fig. 4B**) to normalize the distribution of cell growth over time.

## ACKNOWLEDGEMENTS

We thank Brittany Collins for critical reading of the manuscript and Ozlen Balcioglu and Benjamin Spike for use of their IncuCyte ZOOM cell imager.

## FIGURE LEGENDS

**Table S1.**
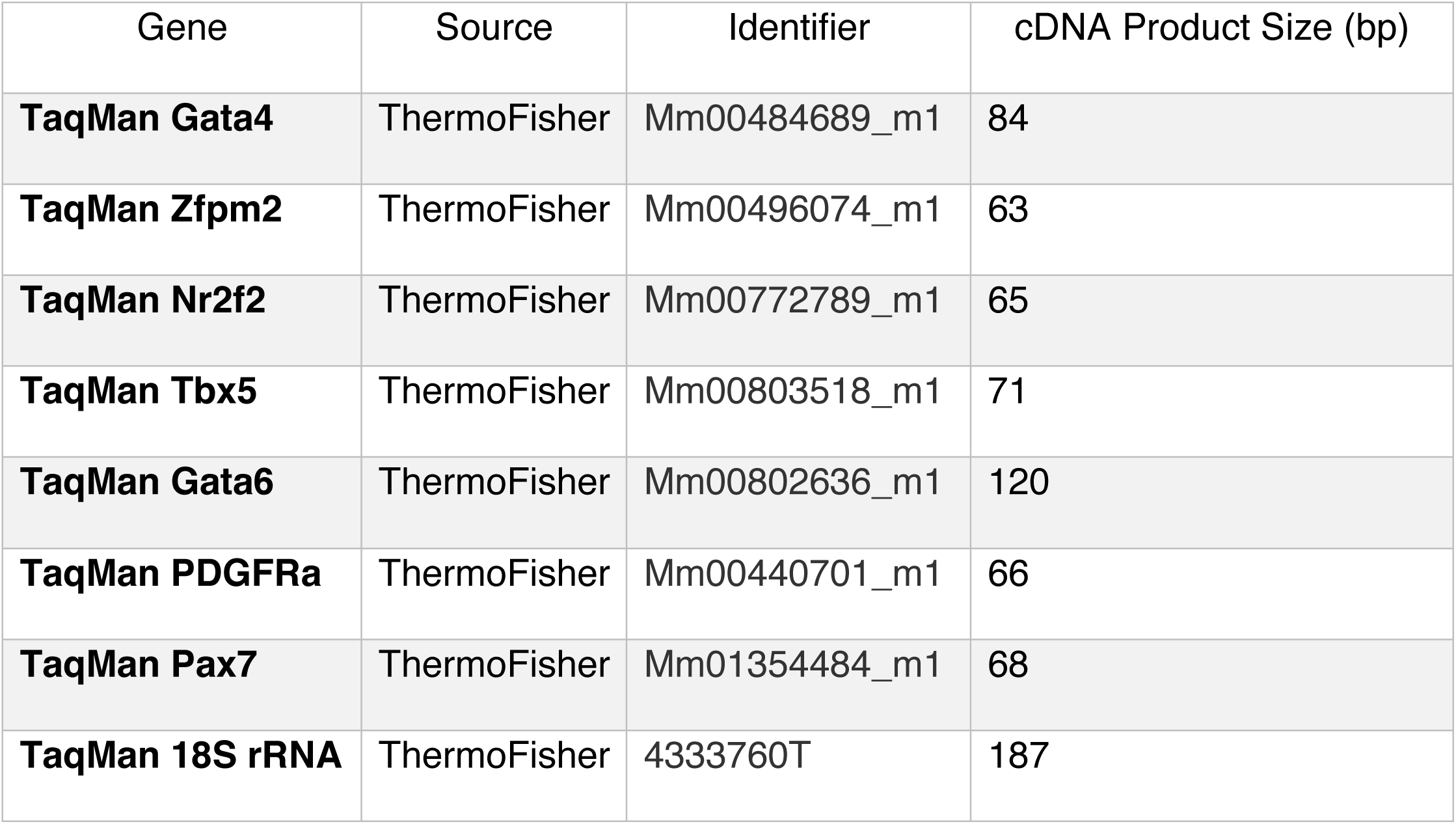
Oligonucleotides for qPCR (related to Figures 3 and Supplemental Figures 1 and 3).

