## Supplementary Figure and Movie Captions for "Cell culture system to efficiently test candidate genes and molecular pathways implicated in congenital diaphragmatic hernias"

**Figure S1. Pax7+ myogenic cells are not maintained in PPFs cultured with PPF growth media. A-B.** Expression of *Pax7* at initial harvest, 5 days of culture, and 10 days of culture of E12.5 PPFs (**A**) and E11.5 PPFs (**B**). Although detected at harvest of E11.5 and E12.5 PPFs, *Pax7* expression is largely undetectable at 5 and 10 days of culture and also absent in NIH3T3 cells. Gene expression determined by qRT-PCR, normalized against *18S* gene expression, and then normalized to "at harvest" value, which was set to 1. Significance tested with one-way ANOVA. Holm–Sidak post hoc tests are indicated by ** $p<0.01$, **** $p<0.0001$. Error bars represent standard error of the mean (SEM).

**Figure S2. Growth of E12.5 PPFs is more variable than NIH 3T3 fibroblasts. A**. Growth of 24 individual wells of PPF or NIH 3T3 fibroblasts. Growth of NIH 3T3 fibroblasts (**B**, n = 9), PPF cells after 5 d in culture and 1 passage (**C**, n = 11), and PPF cells after 10 d in culture and 2 passages (**D**, n = 4). PPFs seeded at lower initial densities generally fail to proliferate (labeled with *'s in **C**, **D**). Growth is measured by percent confluency calculated by IncuCyte software. The average of 4 measurements of confluency +/- SEM is shown for each 2 hour time point.

**Figure S3. Expression of multiple important diaphragm genes is not maintained in E11.5 PPFs during culture.** Expression of *Gata4* (**A**) and *Gata6* (**B**) does not statistically differ in E11.5 PPFs with 5 or 10 d in culture. However, *Zfpm2* (**C**), *Tbx5* (**D**), *Nr2f2* (**E**), and *PDGFRa* (**F**) decrease in E11.5 PPFs in culture. NIH 3T3 fibroblasts have similar levels of *Gata4*, *Gata6*, *Zfpm2*, *Nr2f2*, and *Pdgfr*$\alpha$, but undetectable *Tbx5* as compared with E11.5 PPFs at harvest. (**A**-**F**). **G**. Expression of diaphragm genes does not statistically differ between E11.5 and E12.5 PPFs at harvest. Gene expression determined by qRT-PCR, normalized against *18S* gene expression, and then normalized to "at harvest" value, which was set to 1. Significance tested with one-way ANOVA. Holm–Sidak post hoc tests are indicated by * $p<0.05$, *** $p<0.001$

**** $p<0.0001$. Error bars represent standard error of the mean (SEM).

**Video 1. PPF dissection at E12.5.** Trimmed trunk of embryo with underlying pink liver is pinned on dish with Sylgard. Cranial view of dissection to expose PPFs. Forceps are used to gently pull up left and right sides of the PPFs before completely separating from the midline.

**Video 2. Spreading of the PPFs during the first 4 d in culture.** E11.5 PPFs imaged on IncuCyte ZOOM. PPFs proliferate and expand during initial culture period. Occasional thin and darker myogenic cells are sporadically visible.
