## Supplementary FIgures for "Cell culture system to efficiently test candidate genes and molecular pathways implicated in congenital diaphragmatic hernias"

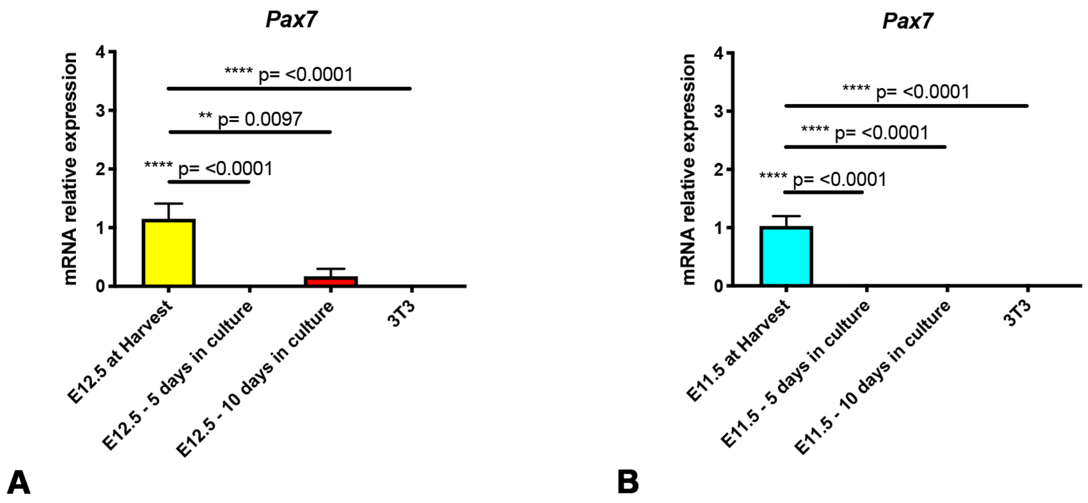

**Bogenschutz, Sefton, and Kardon**
**FIgure S2**

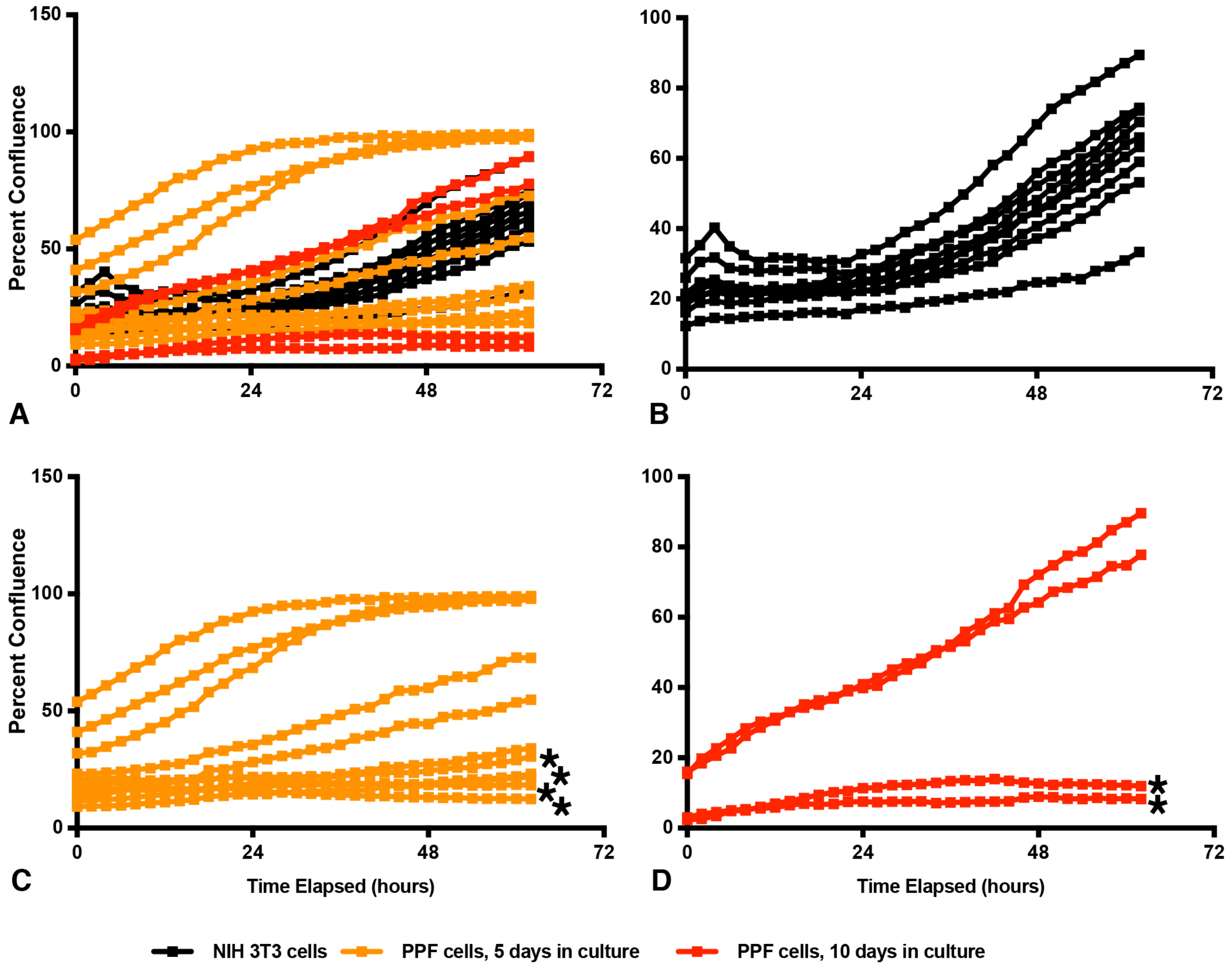

**Bogenschutz, Sefton, and Kardon**
**FIgure S3**

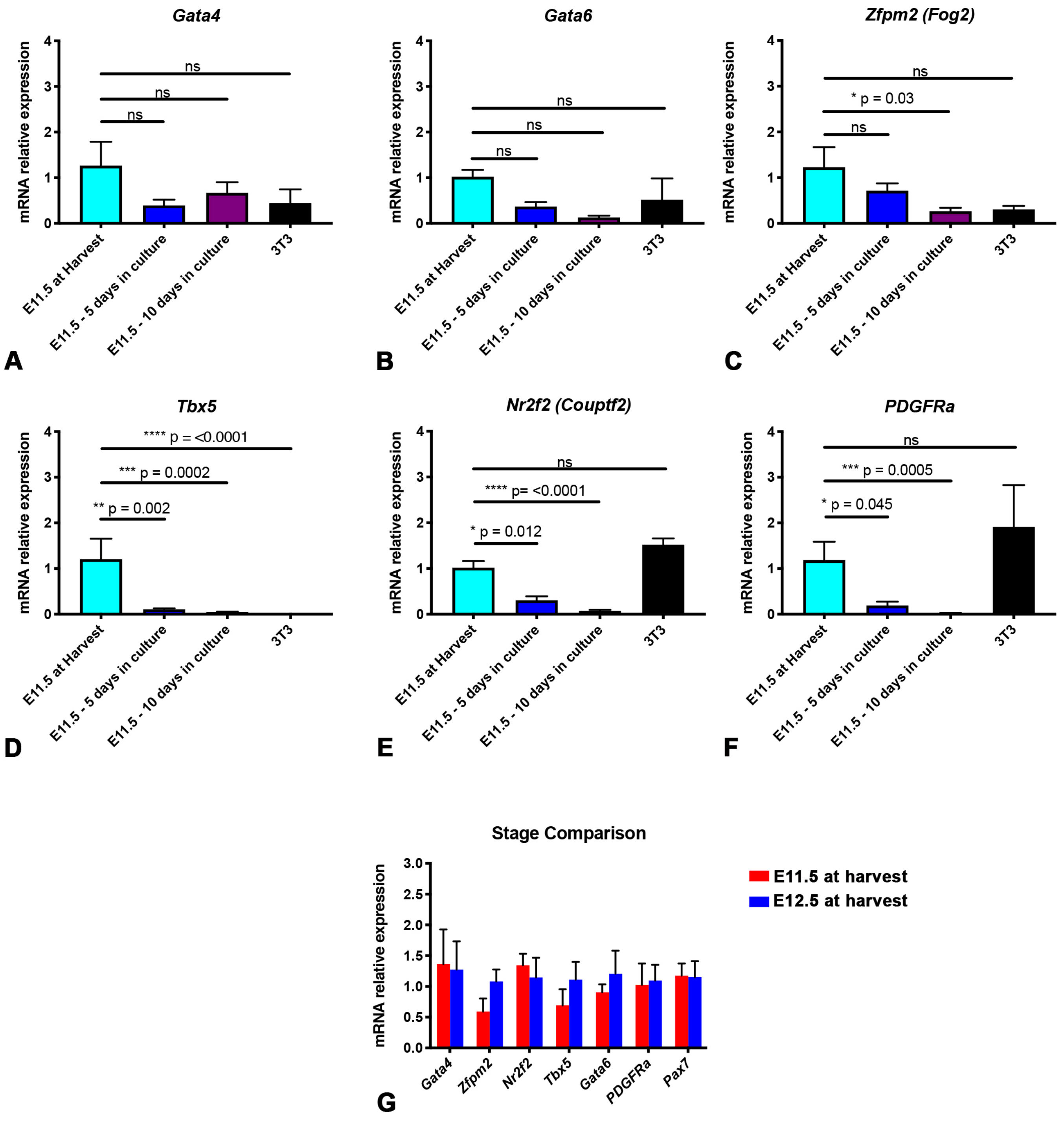
